# Valence biases factual and counterfactual learning in opposite directions

**DOI:** 10.1101/090654

**Authors:** Stefano Palminteri, Germain Lefebvre, Emma J. Kilford, Sarah-Jayne Blakemore

## Abstract

Previous studies suggest that *factual* learning, that is, learning from obtained outcomes, is biased, such that participants preferentially take into account positive, as compared to negative, prediction errors. However, whether or not the prediction error valence also affects *counterfactual* learning, that is, learning from forgone outcomes, is unknown. To address this question, we analysed the performance of two cohorts of participants on reinforcement learning tasks using a computational model that was adapted to test if prediction error valance influences learning. Concerning factual learning, we replicated previous findings of a valence-induced bias, whereby participants learned preferentially from positive, relative to negative, prediction errors. In contrast, for counterfactual learning, we found the opposite valence-induced bias: negative prediction errors were preferentially taken into account relative to positive ones. When considering valence-induced bias in the context of both factual and counterfactual learning, it appears that people tend to preferentially take into account information that confirms their current choice

## Introduction

Goal-directed behaviour is composed of two core components(1): one component is the decisionmaking process that starts from representing the available options and terminates in selecting the option with the highest expected value; the second component is reinforcement learning (RL), through which outcomes are used to refine value expectations in order to improve decision-making. Human decision-making has been shown to be subject to biases (i.e. deviations from the normative prescriptions), such as the framing effect(2). Whereas the investigation of decision-making biases has a long history in economics and psychology, learning biases have been much less systematically investigated(3). This is surprising as most of the decisions we deal with in everyday life are experience-based and choice contexts are often recurrent, thus allowing learning to occur and influence decision-making. In addition, it is important to investigate learning biases as RL processes might play an important role in psychiatric pathogenesis and economic maladaptive behaviour(4, 5).

Standard RL algorithms learn action-outcome associations directly from obtained outcomes on a trial and error basis(6). We refer to this direct form of learning as “factual learning”. Despite the fact that standard models, built around the notion of computational and statistical optimality, prescribe that an agent should learn equally well from positive and negative obtained outcomes (7–9), previous studies have consistently shown that humans display a significant valence-induced bias. The bias generally goes in the direction of preferential learning from positive, compared to negative outcome prediction errors(10–14). This asymmetry in the effects of valence on RL could represent a “low-level” counterpart of the “good news/bad news” effect observed for “high-level” real life prospects, which has been suggested to contribute to maintaining an optimism bias(15).

However, human RL cannot be reduced simply to learning from obtained outcomes. Other sources of information can be successfully integrated in order to improve performance and RL has a multimodular structure(16). Amongst the computational modules that have been demonstrated in humans is counterfactual learning. Counterfactual learning refers to the ability to learn from forgone outcomes (i.e. the outcomes of the option(s) that were not chosen)(17, 18). So far, whether or not a similar valance-induced bias also affects counterfactual remains unknown.

To address this question, we ran two experiments implicating instrumental learning and computational model-based analyses. Two cohorts of healthy volunteers performed variants of a repeated two-armed bandit task involving probabilistic outcomes(19, 20) (**Figure 1A** and **1B**). We analysed the data using a modified version Rescorla-Wagner model that assumes different learning rates for positive and negative, factual and counterfactual, prediction errors (**Figure 1C**)(21, 22).

**Figure 1:**
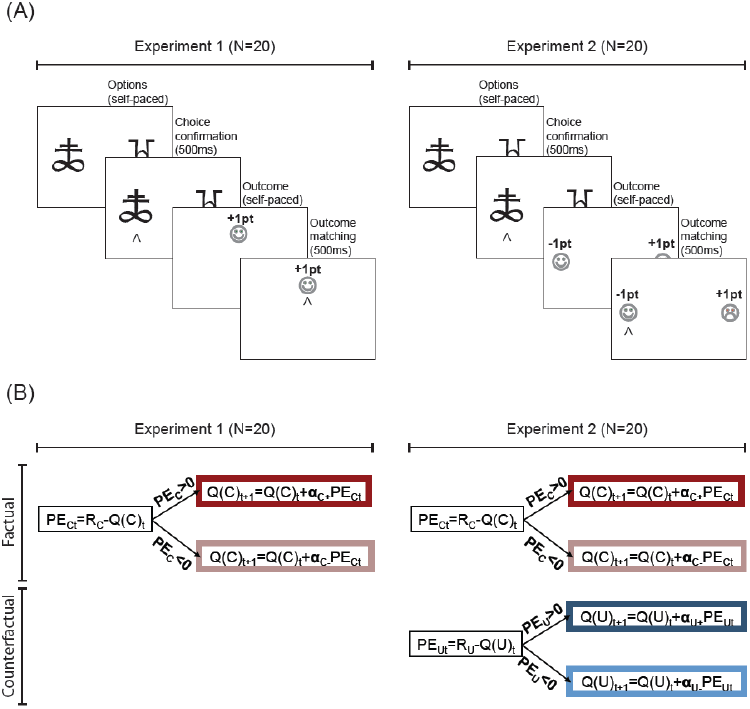
Behavioural task variants and computational model. (**A**) In Experiment 1 (leftmost panel) participants were shown only the outcome of the chosen option. In Experiment 2 (rightmost panel) participants were shown the outcome of both the chosen and the unchosen options. **(B) Computational models**. The schematic summarizes the value update stage of our computational model. The model contains two computational modules, a factual learning module (in red) to learn from chosen outcomes (R_C_) and a counterfactual learning module (in blue) to learn from unchosen outcomes (R_U_) (note that the counterfactual learning module does not apply to Experiment 1). Chosen (Q_C_) and unchosen (Q_U_) option values are updated with delta rules that use different learning rates for positive and negative factual (PE_C_) and counterfactual predictions errors (PE_U_).

The first experiment aimed to replicate previous findings of a “positive valence bias” at the level of factual learning. In this first experiment, participants were presented only with the obtained outcome (chosen outcome: R_C_; **Figure lA**)(10). In the second experiment, in order to investigate whether or not counterfactual learning rates are also affected by the valence of prediction errors, we used a variant of the same instrumental learning task, in which participants were also presented with the forgone outcome (unchosen outcome: R_U_; **Figure 1B**). Our design allowed us to test competing hypotheses concerning the effect of valence on counterfactual learning (**Figure 2A**). A first hypothesis was that, as opposed to factual learning, counterfactual learning would be unbiased. The second hypothesis was that factual and counterfactual learning would present the same valence-induced bias, such that positive counterfactual prediction errors were more likely to be taken into account than negative counterfactual prediction errors. In this scenario the factual and counterfactual learning biases would be consequences of a more general “positive valence” bias, in which positive prediction errors have a greater impact on learning, regardless of whether the option was chosen or not. Finally, the third hypothesis was that valence would affect factual and counterfactual learning in opposing directions, such that negative unchosen prediction errors are more likely to be taken into account than positive unchosen prediction errors. In this last scenario the factual and counterfactual learning biases would be consequences of a more general “confirmation bias”, in which outcomes that support the current choice, are preferentially taken into account.

**Figure 2:**
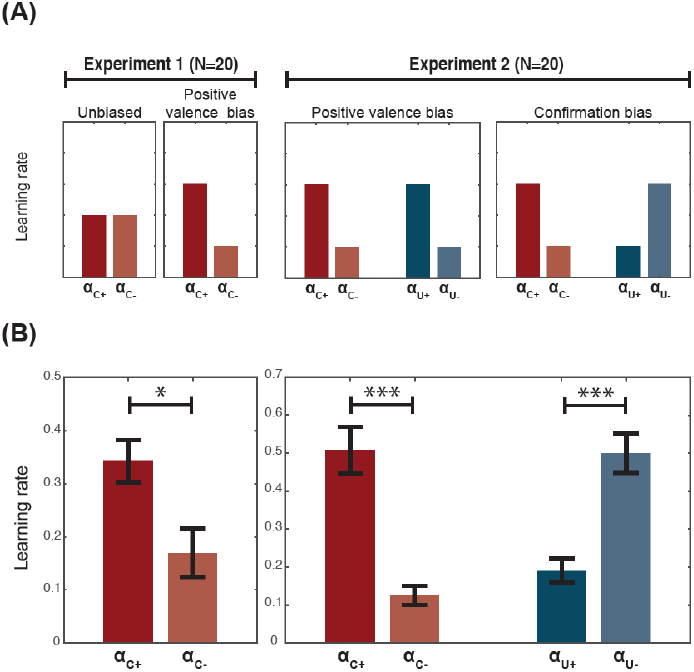
Factual and counterfactual learning biases. (**A**) **Predicted results**. Based on previous studies we expected that in Experiment 1 factual learning would display a “positive valence” bias (i.e. the learning rate for the chosen positive outcomes would be relatively higher than higher than that of the chosen negative outcomes 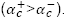 In Experiment 2, one possibility was that this “positive valence” bias would extend to counterfactual learning, whereby positive outcomes are over-weighted regardless of whether the outcome was chosen or unchosen (“valuation” bias) 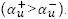. Another possibility was that counterfactual learning would present an opposite bias, whereby the learning rate for the unchosen negative outcomes was higher than the learning rate of the unchosen positive outcomes 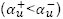 (“confirmation” bias). (**B**) **Actual results**. Learning rate analysis of Experiment 1 replicated previous findings, demonstrating that factual learning presents a “positive valence”, or “optimistic” bias. Learning rate analysis of Experiment 2 indicated that counterfactual learning was also biased, in a direction that was consistent with a “confirmation” bias. \*\*\**P*<0.001 and \**P*<0.05, two-tailed paired t-test.

Learning rate analysis, as a function of both outcome valence (positive and negative) and outcome type (factual and counterfactual), as well as model comparison results, were consistent with this last hypothesis. Behavioural analysis showed that the factual and counterfactual learning biases might be maladaptive, especially in situations involving changes in reward contingencies.

## Results

### Behavioural task and full computational model

To investigate both factual and counterfactual reinforcement learning biases we designed an instrumental task based on a previous design, in which we showed a significant optimistic bias in factual learning(10). The two experiments were different in that the task used in Experiment 1 (N=20) involved participants being shown only the outcome of their chosen option (**Figure 1A**), whereas in Experiment 2 (N=20) the outcome of the unchosen option was also displayed (**Figure 1B**). To test our hypotheses concerning valence-induced learning biases (**Figure 2A**) we fitted the data with a Rescorla-Wagner model assuming different learning rates for positive and negative outcomes, which respectively elicit positive and negative prediction errors (**Figure 1C**). The algorithm used to explain Experiment 1 data involved two learning rates for obtained outcomes (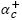 and 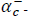 for positive and negative prediction errors of the obtained outcomes, respectively); in addition to the obtained outcomes learning rates, the algorithm used to explain Experiment 2 data also involved two learning rates for forgone outcomes (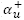 and 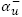 for positive and negative prediction errors of the forgone outcomes, respectively).

### Learning rate analysis

Replicating previous findings, in Experiment 1 we found that the positive factual learning rate (α_C+_) was significantly higher than the negative one (α_C-_; T(19)=2.4; P=0.03) (**Figure 2B**, left). Regarding Experiment 2, we submitted learning rates to a repeated-measure ANOVA with prediction error valence (positive or negative) and prediction error type (factual or counterfactual) as within-subjects factors. Falsifying the “positive valence bias” hypothesis the ANOVA revealed no main effect of prediction error valence (F(1,19)=0.5; P>0.4). We also did not find any effect of outcome type, indicating that, on average, factual and counterfactual learning were similar (F(1,19)=0.2; P>0.6). Consistent with the “confirmation bias” hypothesis we found a significant interaction between valence and type (F(1,19)=119.2; P=1.3e-9). Post-hoc tests indicated that the interaction was driven by effects of valence on both factual (α_C+_>α_C-_; T(19)=6.2; P=6.9e-6) and counterfactual learning rates (α_U-_>α_U+_; T(19)=5.7; P=0.0002) (**Figure 2B** right).

To verify the robustness of this result in the context of different reward contingencies, we analysed learning rates in each task condition separately (**Figure S1A**). We submitted Experiment 1 factual learning rates to a repeated-measure ANOVA with prediction error valence (positive and negative) and task condition as within-subjects factors (**Figure S1B**). Confirming the aggregate result, the ANOVA showed a significant main effect of valence (F(1,19)=26.4, P=5.8e-5), but no effect of condition (F(2,38)=0.7, P>0.5) and, crucially, no valence by condition interaction (F(2,38)=0.8, P>0.4). We submitted Experiment 2 factual and counterfactual learning rates to a repeated-measure ANOVA with prediction error valence (positive and negative), prediction error type (factual or counterfactual) and condition (Symmetric, Asymmetric and Reversal) as within-subjects factors (**Figure S1C**). Confirming the aggregate result, the ANOVA showed no effect of valence (F(1,19)=0.0, P>0.9), no effect of type (F(1,19)=0.3, P>0.5), but a significant valence by type interaction (F(1,19)=162.9, P=9.1e-11). We also found an effect of condition (F(2,38)=5.1, P=0.01), reflecting lower average learning rates in the Reversal compared to the Asymmetric condition (T(19)=2.99; P=0.007), which was not modulated by valence (F(2,38)=0.2, P>0.7), or type (F(2,38)=1.2, P>0.3). Crucially, the three-way interaction was not significant (F(2,38)=1.8, P>.1). These results indicate that learning biases were robust across different task contingencies.

### Dimensionality reduction with model comparison

To further test our hypotheses and verify its parsimony, we ran a model comparison analysis including the ‘Full’ model (i.e., the model with four learning rates; **Figure 1C**) and reduced versions of it (**Figure 3A**). The first alternative model was obtained by reducing the number of learning rates along the dimension of the outcome type (factional or counterfactual). This ‘Information’ model has only two learning rates: one for the obtained outcomes (α_C_) and another for the forgone outcomes (α_U_). The second alternative model was obtained by reducing the number of learning rates along the dimension of the outcome valence (positive or negative). This ‘Valence’ model has only two learning rates (one for the positive outcomes (α_+_) and another for the negative outcomes (α_-_)) and should win according to the “positive valence bias” hypothesis. Finally, the third alternative model was obtained by reducing the learning rate as a function of the outcome event being confirmatory (positive obtained or negative forgone) or disconfirmatory (negative obtained and positive forgone). This ‘Confirmation’ model has only two learning rates (one for confirmatory outcomes (α_CON_) and another for the disconfirmatory outcomes (α_DIS_)) and should win according to the “confirmation bias” hypothesis.

**Figure 3:**
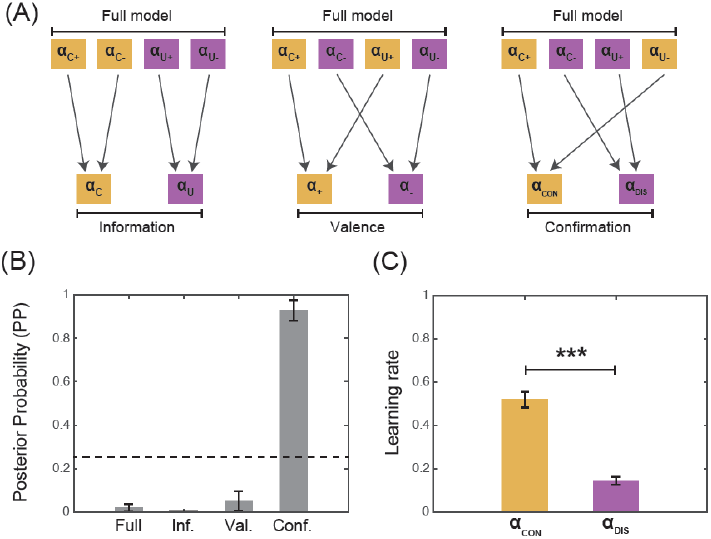
dimensionality reduction with model comparison. **(A**) **Model space**. The figure represents how the number of parameters (learning rate) is reduced moving from the ‘Full’ model to more simple ones. (**B**) **Model Comparison**. The panel represents the posterior probability (PP) of the models, whose calculation is based on the BIC that penalizes model complexity. The dashed line represents random posterior probability (0.25). (C) Model parameters. The panel represents the learning rate for the best fitting model (i.e., the ‘Confirmation’) model: αCON: learning rate for positive obtained and negative forgone outcomes; αDIS: learning rate for negative obtained and positive forgone outcomes. \*\*\**P*<0.001, two-tailed paired t-test.

**Figure 4:**
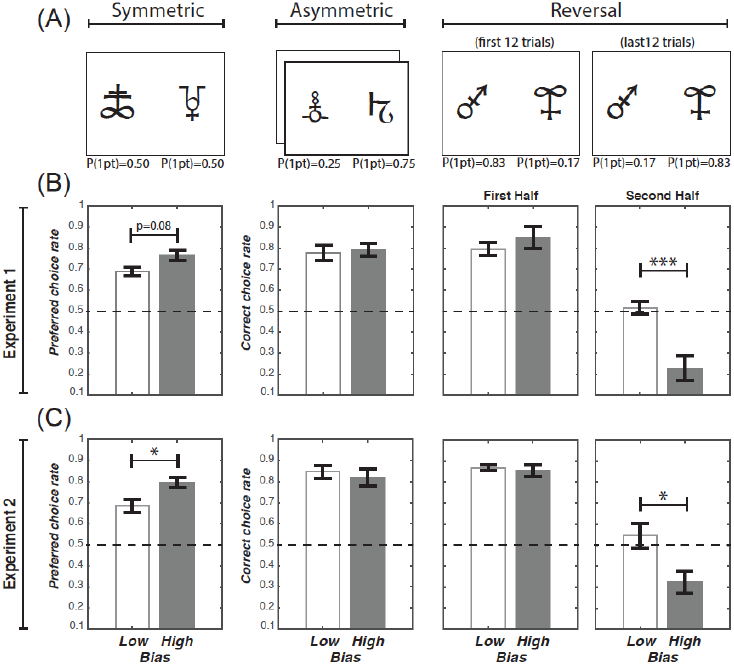
Behavioral signatures distinguishing “low” and “high bias” participants. (**A**) **Task conditions**. The ‘Symmetric’ condition was characterized by a stable reward contingency and no correct option, because the two options have equal reward probabilities. The ‘Asymmetric conditions’ were also characterized by a stable reward contingency and a correct option, since one option had a higher reward probability than the other. The ‘Reversal’ condition was characterized by an instable reward contingency: after 12 trials the reward probability reversed across symbols, so that the former correct option became the incorrect one, and vice versa. Note that the number of trials refers to one session and participants performed two sessions, each involving new pairs of stimuli (192 trials in total). (**B**) and (**C**) Behavioural results as a function of the task conditions in Experiment 1 and Experiment 2, respectively. Each column presents the result of the corresponding condition presented in (**A**). In the Symmetric conditions, where there was no correct option, we calculated the “Preferred choice rate”, which was the choice rate of the most frequently chosen option (by definition, this was always greater than 0.5). In the Asymmetric and the Reversal conditions we calculated the correct choice rate. In the Reversal condition the correct choice rate was split between the two learning phases. \*\*\**P*<0.001 and \**P*<0.05, two-tailed paired t-test.

BIC analysis indicated that the ‘Full’ model better accounted for the data compared to both the ‘Information’ and the ‘Valence’ models (both comparisons: T(19)>4.2; P<0.0005; **Table 1**). However the ‘Confirmation’ model better accounted for the data compared to the ‘Full’ model (T(19)=9.9; P=6.4e-9). The posterior probability of the ‘Confirmation’ model was higher compared to the chance (.0.25 for a model space including 4 models; T(19)=13.5; P= 3.3e-11) and compared to that of all the other models (all comparison: T(19)>9.0; P<2.1e-8) and the learning rate for confirmative outcomes was significantly higher compared that for disconfirmatory events (α_CON_>α_DIS_; T(19)=11.7; P=3.9e-10) (**Figure 3B and Figure 3C**). These results support the “confirmation bias” hypothesis and further indicate that, at least at the behavioural level, chosen and unchosen outcomes may be processed by the same learning systems.

**Table 1:** Model Comparison. BIC: Bayesian Information Criterion; PP: posterior probability; XP: exceedance probability; df: degrees of freedom. The “winning” model is the “Confirmation”, whose learning rates are displayed in **Figure 3C**.The second best model is the “Full” model, whose learning rates are displayed in **Figure 2C**.

| Model | Full (5df) | Information (3df) | Valence (3df) | Confirmation (3df) | Perseveration (4df) | One (2df) |
| --- | --- | --- | --- | --- | --- | --- |
| BIC | 162.0±13.4 | 178.2±13.0 | 180.7±11.8 | 155.0±13.2 | 165.2±13.6 | 179.1±11.8 |
| PP | 0.02±0.02 | 0.00±0.00 | 0.05±0.05 | 0.89±0.06 | 0.01±0.01 | 0.04±0.03 |
| XP | 0.0 | 0.0 | 0.0 | 1.0 | 0.0 | 0.0 |

### Behavioural signatures of learning biases

To investigate the behavioural consequences of the learning biases we median-split all participants according their normalised learning rate differences. We reasoned that the effects of learning biases on behavioural performance could be highlighted comparing participants who differed in the extent they expressed the bias itself. Experiment 1 participants were split according to their normalised factual learning rate bias: 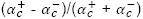, from which we obtained a high (0.76±0.05) and a low bias (0.11±0.14) group. Experiment 2 participants were split according their normalised confirmation learning rate bias: 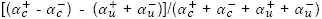, from which we also obtained a high (0.72±0.04) and a low bias (0.36±0.04) group.

In both experiments, our task included three different conditions (**Figure S1A**): a “Symmetric” condition, in which both options were associated with a 50% chance of getting a reward; an “Asymmetric” condition, in which one option was associated with a 75% chance of getting a reward, whereas the other option was associated with only a 25% chance; a “Reversal” condition, in which one option was initially associated with a 83% chance of getting a reward and the other option was associated with a 17% chance of getting a reward, but after 12 trials the contingency reversed. From the Symmetric condition we extracted preferred choice rate as a dependent variable, which was the choice rate of the most frequently chosen option (i.e. the option/symbol that was chosen more than >50%). We submitted the preferred choice rate to an ANOVA with experiment (1 or 2) and bias level (high and low) as between-subjects factors. The ANOVA showed a significant main effect of bias level (F(1,36)=8.8, P=0.006). There was no significant main effect of experiment (F(1,36)=0.6, P>0.6) and no significant interaction between experiment and bias level (F(1,36)=0.3, P>0.5). The main effect of bias level was driven by higher preferred choice rate in the high, compared to the low bias group in both Experiment 1 (T(18)=1.8 P=0.08) and Experiment 2 (T(18)=2.3 P=0.03). This result suggests that higher biases were associated with an increased tendency to develop a preferred choice, even in the absence of a “correct” option, which naturally emerges from overweighting positive outcomes(10).

From the remaining conditions we extracted the correct choice rate, which was the choice rate of the most frequently rewarded option. In the Reversal condition, correct choice rate was split across the first (i.e., before the reversal of the contingencies) and second half (i.e., after the reversal of the contingencies) of the trial. We submitted the correct choice rate to a mixed ANOVA with experiment (1 or 2) and Bias Group (high and low) as between-subjects factors, and condition (Asymmetric, Reversal: first half, and Reversal: second half) as a within-subjects factor. We found a main effect of experiment (F(1,36)=4.1, P=0.05), indicating that correct choice rate was higher in Experiment 2 than Experiment 1, which is consistent with previous studies showing that counterfactual feedback enhances learning(20, 23). We also found a significant effect of bias level (F(1,36)=10.8, P=0.002), a significant effect of condition (F(2,72)=99.5, P=2.0e-16), and a significant bias level by condition interaction (F(2,72)=9.6, P=0.0002). Indeed, in both experiments, the correct choice rate in the second half of the Reversal condition was lower in the high bias compared to the low bias group (Experiment 1: T(18)=3.9 P=0.0003; Experiment 2: T(18)=2.5 P=0.02). This result derives from the fact that in the first half of the Reversal condition learning is primarily driven by positive factual prediction errors (and negative counterfactual prediction errors, where applicable), whereas in the second half of the Reversal condition correct performance depends on un-learning previous associations, based on negative factual prediction errors (and positive counterfactual prediction errors, in Experiment 2).

## Discussion

Two cohorts of healthy adult participants performed two variants of an instrumental learning task, involving factual (Experiment 1) and counterfactual (Experiments 1 & 2) reinforcement learning. We found that prediction error valence biased factual and counterfactual learning in opposite directions. When learning from obtained outcomes (factual learning), the learning rate for positive prediction errors was higher than the learning rate for negative prediction errors. When learning from forgone outcomes (counterfactual learning), the learning rate for positive prediction errors was lower than that of negative prediction errors. This result proved stable across different reward contingency conditions and was further supported by model comparison indicating that the most parsimonious model was a model with different learning rates for confirmatory and disconfirmatory events, regardless of outcome type (factual or counterfactual) and valence (positive or negative). Finally, model-free analysis showed that participants with a higher valence-induced learning bias displayed poorer learning performance, specifically when it was necessary to adjust their behaviour in response to a reversal of reward contingencies.

Our results demonstrating a factual learning bias replicate previous findings showing that in simple instrumental learning tasks, participants preferentially learn from positive compared to negative prediction errors(11–13). However, in contrast to previous studies in which this learning bias had no negative impact on behavioural performance (i.e., correct choice rate and therefore final payoff), here we demonstrated that this learning bias is still present in situations where it has a negative impact on performance. In fact, whereas low and high bias participants performed equally well in conditions with stable reward contingencies, in conditions with unstable reward contingencies we found that high bias participants showed a relatively reduced correct choice rate. In the Reversal condition, learning to successfully reverse the response in the second half of the trials is mainly driven by negative factual (and positive counterfactual) prediction errors, however participants displaying higher biases presented a lower correct choice rate, specifically in the second half of the “Reversal” condition.

In addition to reduced reversal learning, and in accordance with a previous study(10), another behavioural feature that distinguished higher and lower bias participants was the preferred response rate in the Symmetric condition. In the Symmetric condition, both cues had the same reward probabilities (50%), such that there was no intrinsic “correct” response, allowing us to calculate a “preferred” response rate for each participant (defined as the choice rate of the option most frequently selected by a given participant, i.e. the option selected in > 50% of trials). The preferred response rate can therefore be taken as a measure of the tendency to overestimate the value of one cue compared to the other, in the absence of actual outcome-based, factual evidence. In both experiments, higher bias participants showed higher preferred response rates, a behavioural pattern that is consistent with an increased tendency to discount negative factual (and positive counterfactual) prediction errors, which can result in one considering a previously rewarded chosen option as better than it really is and an increased preference for this choice.

Previous studies have so far been unable to distinguish whether this valence-induced factual learning bias was a valuation or a confirmation bias. In other words, do participants preferentially learn from positive prediction errors because they are positively valenced or because the outcome “confirms” the choice they have just made? To address this question we designed Experiment 2, where, by the inclusion of counterfactual feedback, we were able to separate the influence of valence (positive vs. negative) from the influence of choice (chosen vs. unchosen). Crucially, whereas the two competing hypotheses (“valuation” vs. “confirmation”) predict the same result concerning factual leaning rates, they predict opposite effects of valence on counterfactual learning rates. The results from Experiment 2 support the “confirmation” bias hypothesis: participants preferentially took into account the outcomes that “confirmed” their current behavioural policy (positive chosen and negative unchosen outcomes) and discounted the outcomes that contradicted it (negative chosen and positive unchosen outcomes). Our results therefore support the idea that confirmation biases are pervasive in human cognition(24).

It should be noted that, from an orthodox Bayesian perspective, a “confirmation bias” would involve reinforcing one’s own initial beliefs or preferences. Previous studies have investigated how prior information — in the form of explicit task instructions or advice — influences the learning of reinforcement statistics and have provided evidence of a confirmation bias(25–27). However, consistent with our study, their computational and neural results suggest that this instruction-induced confirmation bias operates at the level of outcome processing and not at the level of initial preferences nor at the level of the decision-making process(28, 29). Here, we take a slightly different perspective by extending the notion of confirmation to the implicit reinforcement of one’s own current choice, independently from explicit prior information, by preferentially learning from desirable outcomes.

We performed our learning rate analysis separately for each task condition and the results proved robust and not driven by any particular reward contingency condition. While our results contrast with previous studies that have found learning rates adapted as a function of task contingencies, showing increases when task contingencies were unstable(30, 31), several differences between these tasks and ours may explain this discrepancy. First, in previous studies the stable and unstable phases were clearly separated, whereas in our design participants were simultaneously tested with the three reward contingency conditions. Second, in our experiments we did not explicitly tell participants to monitor the stability of the reward contingency. Finally, since in our task the Reversal condition represented only fourth quarter of the trials, participants may not have explicitly realized that changing learning rates were adaptive in some cases.

To date, two different views of counterfactual learning have been proposed in the literature. In one view, factual and counterfactual learning are underpinned by different systems that could be computationally and anatomically mapped into subcortical - model-free - and prefrontal - model-based - modules(17, 18, 32). In another view, factual and counterfactual outcomes are proposed to be processed by the same learning system, involving the dopaminergic nuclei and their projections(33–35). Our dimensionality reduction model comparison result sheds new light on this debate. According to the first view that factual and counterfactual learning are based on different systems, different learning rates for positive and negative prediction errors would have better accounted the data (the ‘Information’ model). On the contrary, the winning model assumes the learning process is different across desirable and undesirable outcomes but shared across obtained and forgone outcomes (the ‘Confirmation’ model), which supports the view that factual and counterfactual learning are different facets of the same system.

Why do these learning biases have an overall maladaptive value? One possibility is that these learning biases are maladaptive, but they arise from neurobiological constraints, which limit human learning capacity. However, we believe this interpretation is unlikely because we see no clear reason why such limits would differentially affect learning from positive and negative prediction errors. In other words, we would predict that a neurobiological constraint on learning rate would limit all learning rates in a similar way and therefore not produce valence-induced learning asymmetries.

A second possibility is that these learning biases are not maladaptive. For instance it has been shown that in certain reward conditions agents displaying valence-induced learning bias may outperform unbiased agents (9). Thus, a possible explanation for these learning biases is that they have been positively selected because they can be adaptive in the context of the natural environment in which the learning system evolved(36).

Finally, a third, intermediate possibility is that these learning biases can be maladaptive in the context of learning performance, but due to their adaptive effects in other domains of cognition, overall they have a net adaptive value. For example, these biases may also manifest as “self-serving”, choice-supportive biases, which result in individuals tending to ascribe success to their own abilities and efforts, but relatively tending to neglect failures(37). These psychological processes may help nourish self-esteem and confidence, both of which have been associated with overall favourable real life outcomes(38).

To conclude, by showing that both factual and counterfactual learning are affected by valence-induced biases, our study highlights the importance of investigating reinforcement learning biases. Most of the decisions we face everyday are experience-based, therefore increasing our understanding of learning biases will likely enable the refinement of existing models of value-based decision-making, furthering our understanding of human cognition(3).

## Methods

### Participants

The study included two experiments. Each experiment involved N=20 participants (Experiment 1: 7 males, mean age 23.9 ± 0.7; Experiment 2: 4 males, mean age 22.8 ± 0.7). The local ethics committee approved the study. All participants gave written informed consent before inclusion in the study, which was carried out in accordance with the declaration of Helsinki (1964, revised 2013). The inclusion criteria were being older than 18 years and reporting no history of neurological or psychiatric disorders.

### Behavioural tasks

Participants performed a probabilistic instrumental learning task based on previous studies(19, 20) (Fig. 1A & 1B, Fig. 3A). Briefly, the task involved choosing between two cues that were presented in fixed pairs and therefore represented fixed choice contexts. Cues were associated with stationary outcome probability in three out of four contexts. In the remaining context outcome probability was non-stationary. The possible outcomes were either winning or losing a point. To allow learning, each context was presented 24 trials. Each session comprised the four learning contexts and therefore included 96 trials. The whole experiment involved two sessions, each including the same number of contexts and conditions, but a different set of stimuli. Thus, the total experiment included 192 trials. The four learning contexts (i.e. fixed pairs of cues) were divided in three conditions. In the “Symmetric” condition each cue was associated with a .50 probability of winning one point. In the “Asymmetric” condition one cue was associated with a .75 probability of winning a point and the other cue was associated with a .25 probability of winning a point. The Asymmetric condition was implemented in two choice contexts in each session. Finally, in the “Reversal” condition one cue was associated with a .83 probability of winning a point and the other cue was associated with a .17 probability of winning a point, during the first 12 trials, and these contingencies were reversed thereafter. We chose a bigger probability difference in the Reversal compared to the Asymmetric condition in order to ensure that participants were able to reach a plateau within the first 12 trials. Participants were encouraged to accumulate as many points as possible and were informed that some cues would result in winning more often than others. Participants were given no explicit information regarding reward probabilities, which they had to learn through trial and error.

At each trial, after a fixation cross, the choice context was presented. Participants made their choice by pressing left or right arrow keys with their right hand (the choice time was self-paced). The two experiments differed in the fact that in the Experiment 1 participants were only informed about the outcome of their own choice (chosen outcome), whereas in the Experiment 2 participants were informed about both the obtained and the forgone outcome (i.e. counterfactual feedback). In Experiment 1 positive outcomes were presented on the top and negative outcome of the bottom of the screen. The participant was required to press the key corresponding to the position of the outcome on the screen in order to move to the subsequent trial (top/bottom). In Experiment 2 the obtained outcomes were presented in the same place of the chosen cue and the forgone outcome in the same place of the unchosen cue. To move to the subsequent trial, participants had to match the position of the outcome (right/left). Importantly for our computational analyses, outcome probabilities (although on average anti-correlated in the Asymmetric” and Reversal conditions) were truly independent across cues, so that in the Symmetric condition, in a given trial, the obtained and forgone outcomes were the same in the 50% of the trials; in the Asymmetric condition this was the case in the 37.5% of the trials; finally, in the Reversal condition this was the case in the 28.2% of the trials.

### Behavioural variables

We extracted the correct response rate, that is, the rate of the trials in which the participants chose the most rewarding response, from the Asymmetric and the Reversal conditions. The correct response rate in the Reversal condition was calculated separately for the two phases: before (“first half”) and after (“second half”) the contingency reversal. In the Symmetric condition, we calculated the so-called “preferred” response rate. The preferred response was defined as the most frequently chosen option, i.e. that chosen by the participant on more than 50% of the trials. This quantity is therefore, by definition, greater than 0.5.

### Computational models

We fitted the data with a standard Q-learning model, including different learning rates following positive and negative prediction errors and containing two different modules (Fig 1.C): a factual learning module to learn from chosen outcomes (R_c_) and a counterfactual learning module to learn from unchosen outcomes (R_u_) (note that counterfactual learning applies only to Experiment 2). For each pair of cues (choice context), the model estimates the expected values of the two options (Q-values). These Q-values essentially represent the expected reward obtained by choosing a particular option in a given context. In both experiments, Q-values were set at 0 before learning, corresponding to the a priori expectation of 50% chance of winning 1 point, plus a 50% chance of losing 1 point. After every trial *t*, the value of the chosen option is updated according to the following rule (factual learning module):

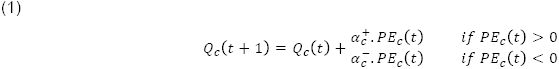

In this first equation, *PE_c_*(*t*) is the prediction error of the chosen option, calculated as:

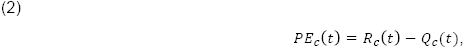

where *R_c_*(*t*) was the reward obtained as an outcome of choosing *c* at trial *t*. In other words, the prediction error *PE_c_*(*t*) is the difference between the expected outcome *Q_c_*(*t*) and the actual outcome *R_c_*(*t*).

In Experiment 2 the unchosen option value was also updated according to following rule (counterfactual learning module):

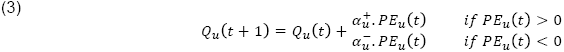

In this second equation, *PE_u_*(*t*) is the prediction error of the unchosen option, calculated as:

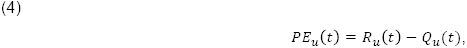
where *R_u_*(*t*) was the reward that could have been obtained as an outcome of having chosen *u* at trial *t*. In other words, the prediction error *PE_u_*(*t*)is the difference between the expected outcome *Q_u_*(*t*) and the actual outcome *R_u_*(*t*) of the unchosen option.

The learning rates 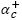 and 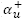 are scaling parameters that adjust the amplitude of value changes from one trial to the next when prediction errors of chosen and unchosen option respectively are positive (when the actual reward *R*(*t*) is better than the expected reward *Q*(*t*)) and the learning rates 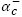 and 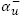 do the same when prediction errors are negative. Thus, our model allows for the amplitude of the update to be different following positive and negative prediction errors and for both chosen and unchosen options. It therefore allows for the existence of valence-dependent learning biases.

Finally, the probability (or likelihood) of selecting the chosen option was estimated with a the soft-max rule as follow:

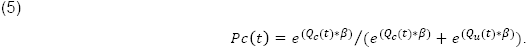

This is a standard stochastic decision rule that calculates the probability of selecting one of a set of options according to their associated values. The temperature, *β*, is another scaling parameter that adjusts the stochasticity of decision-making.

In addition to this ‘Full’ model, we also considered versions with a reduced number of learning rates (**Figure 3A**): the ‘Information’ model, where 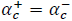 and 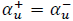; the ‘Valence’ model, where 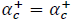 and 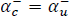; and the ‘Confirmation’ model, where 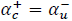 and 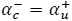. For the model comparison, we also considered a very simple model (the ‘One’) model, with only one learning rate 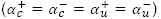, and a model where an additional parameter -Inf < π < +Inf biases the decisionmaking process by increasing (positive values) or decreasing (negative values) the likelihood of repeating the same choice, regardless of the previous outcome (Table 1).

### Parameter optimization and model comparison

In a first analysis, we optimized model parameters by minimizing the negative log-likelihood of the data, given different parameter settings, using Matlab’s fmincon function (ranges: 0<*β*<Infinite, and 0< *α_n_*<1):

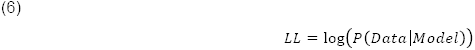

Negative log-likelihoods (LL) were used to compute at the individual level (random effects) for each model the Bayesian information criterion (BIC), as follows:

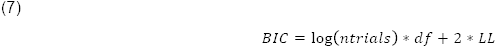

BIC where compared between biased and unbiased models to verify that the utilization of the biased model is justified, even accounting for its extra-complexity. As an approximation of the model evidence, individual BICs were fed into the mbb-vb-toolbox(39), a procedure that estimates the expected frequencies and the exceedance probability for each model within a set of models, given the data gathered from all participants. Expected frequency is a quantification of the posterior probability of the model (denoted PP), i.e. the probability of the model generating the data obtained from any randomly selected participant. Exceedance probability (denoted XP) is the probability that a given model fits the data better than all other models in the set, i.e. has the highest PP (**Table 1**).

In a second analysis, we optimized model parameters by minimizing the logarithm of the Laplace approximation to the model evidence (or log posterior probability: LPP):

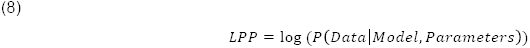

Because LPP maximization includes priors over the parameters (temperature: gamma(1.2,5); learning rates beta(1.1,1.1)) (REF), it avoids degenerate parameter, estimates. Therefore, learning rates analyses have been performed on the values retrieved with this procedure. To avoid bias in learning rate comparison, the same priors were used for all learning rates. In the main analysis, a single set of parameters was used to fit all conditions. In a control analysis, different sets of parameters were used to fit each condition (“Symmetric”, “Asymmetric” and “Reversal”).

### Parameter recovery

To validate our results, and more specifically to verify that valence-induced differences in learning rates reflect true differences in learning, as opposed to an artefact of the parameter optimization procedure, we checked the capacity of recovering the correct parameters in simulated datasets. To do so, we simulated performance on our behavioural task by virtual participants with different learning rates (**Fig. S2 & Fig. S3**). Concerning Experiment 1, we simulated unbiased 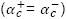 and biased 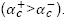 participants. Concerning Experiment 2, we simulated unbiased (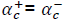 and 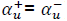), semi-biased (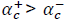 and 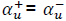) and biased (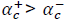 and 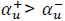) participants. We simulated N=100 virtual participants per set of parameter. The results of these analyses are presented in the supplementary materials and confirm the capacity of our parameter optimization procedure to correctly recover the true parameters, regardless to presence (or absence) of biases (**Fig. S2** and **Fig. S3**).

---

**Figure S1:**
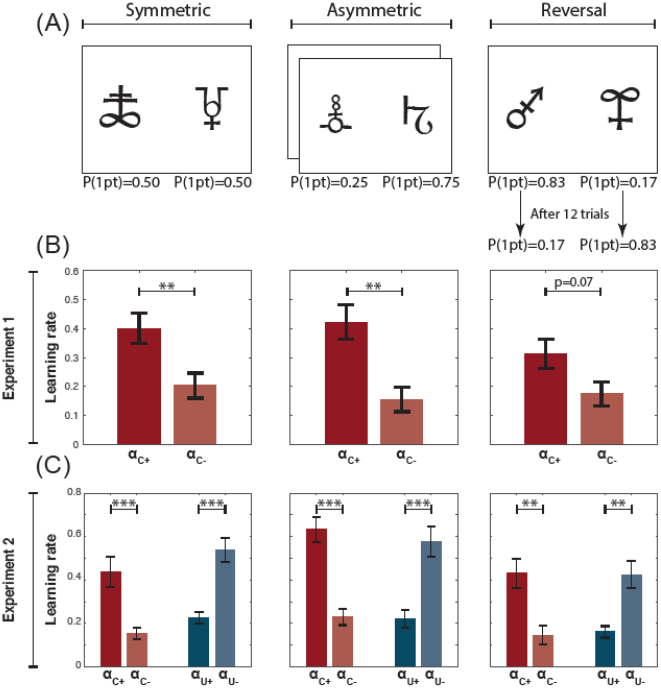
Stability of learning biases across task conditions. (**A**) **Task conditions**. The ‘Symmetric’ condition was characterized by a stable reward contingency and no correct option, because the two options have equal reward probabilities. The ‘Asymmetric conditions’ were also characterized by a stable reward contingency and a correct option, since one option had a higher reward probability than the other. The ‘Reversal’ condition was characterized by an instable reward contingency: after 12 trials the reward probability reversed across symbols, so that the former correct option became the incorrect one, and vice versa. Note that the number of trials refers to one session and participants performed two sessions, each involving new pairs of stimuli (192 trials in total). (**B**) and (**C**) Computational results as a function of the task conditions in Experiment 1 and Experiment 2, respectively. Each column presents the result of the corresponding condition presented in (**A**). \*\*\**P*<0.001 and \*\**P*<0.01, two-tailed paired t-test.

**Supplementary Figure 2:**
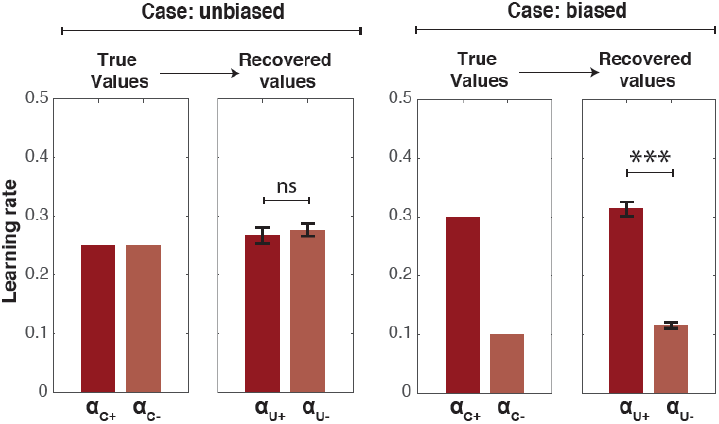
Parameter recovery in the Experiment 1 setting. “True values”: learning rates used to simulate the data. “Recovered values”: learning rates obtained from the simulations once the same parameter optimization was applied as for the experimental data. “Case: unbiased”: no learning rate bias. “Case: biased”: optimistic learning rate bias.

**Supplementary Figure 3:**
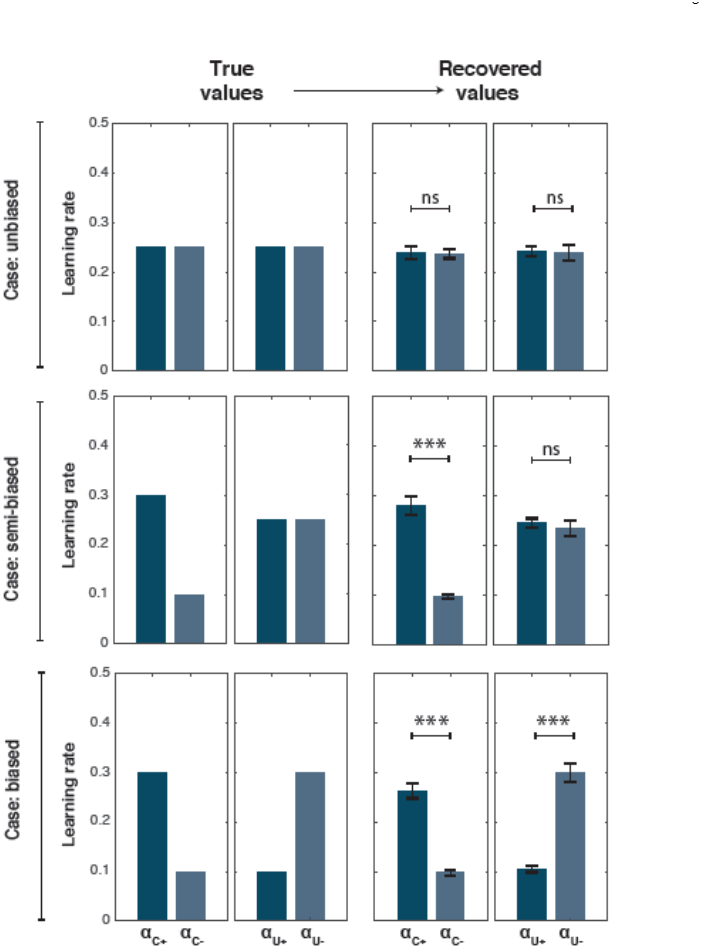
parameter recovery in the Experiment 2 setting. “True values”: learning rates used to simulate the data. “Recovered values”: learning rates obtained from the simulations once applied the same parameter optimization as for the experimental data. “Case: unbiased”: no learning rate bias. “Case: semi-biased”: learning rate bias only concerning factual learning. “Case biased”: confirmation bias involving both factual and counterfactual learning.

## Acknowledgments

Anahit Mkrtchian and Anders Jespersen performed the experiments. We thank Bahador Bahrami and Valerian Chambon for helpful discussions. We thank Vasilisa Skvortsova for a discussion leading to the dimensionality reduction analysis.

